# Carrion converging: Skull shape is predicted by feeding ecology in vultures

**DOI:** 10.1101/2023.05.17.541171

**Authors:** Katherine R Steinfield, Ryan N Felice, Mackenzie E Kirchner, Andrew Knapp

## Abstract

The link between skull shape and dietary ecology in birds at macroevolutionary scales has recently been called into question by analyses of 3D shape that reveal that cranial anatomy is mainly influenced by other factors such as allometry. It is still unknown whether this form-function disconnect also exists at smaller evolutionary scales, for example within specialized ecological guilds. Vultures are a diverse guild of 23 extant species in two families (Accipitridae and Cathartidae) that exhibit phenotypic convergence as a result of highly-specialized feeding ecology. Vultures are the only known obligate scavengers among vertebrates and are usually grouped together under this single dietary category, but within this specialized diet there are three distinct, species-specific feeding strategies termed ripper, gulper, and scrapper. We use three-dimensional geometric morphometrics to quantify the relative contributions of feeding ecology, allometry, and phylogeny on vulture skull shape, along with several non-vulture raptors of similar size, range and ecology. Families show clear separation in shape, but phylogenetic signal is comparatively weak (*K_mult_* = 0.33). Taking into account the influence of phylogeny, skull shape is not significantly correlated with either skull size or feeding type, but there are examples of strong, significant convergence and parallel shape evolution across feeding groups. Furthermore, skull shape performs strongly in predicting feeding ecology in a phylogenetic discriminant function analysis. These findings highlight the importance of detailed assessment of feeding behavior in studies of ecomorphology, rather than broader dietary categories alone, and reveal that ecology can be readily inferred from form given appropriate information.

## Introduction

The avian skull has long been an exemplar of adaptive evolution due to the incredible phenotypic diversity in extant birds, bringing to mind the most classic examples such as Darwin’s finches or Hawaiian Honeycreepers (Pigot et al., 2016; Cooney et al., 2017; Lack, 1953; Smith et al., 1995; Lovette et al., 2002; Grant & Grant, 2006; Gibbs & Grant, 1987; Olsen, 2017; Jønsson et al., 2012; Felice et al., 2019). Attempts to quantify this classic association between form and function through three-dimensional geometric morphometrics have demonstrated that diet insufficiently explains the majority of shape variation, with factors such as allometry or phylogeny contributing significantly more (Felice et al., 2019; Bright et al., 2016; Navalon et al., 2019; Bright et al., 2019). Within more restricted taxonomic groups, geometric morphometric analyses have found that diet has low explanatory power (Bright et al., 2019), whereas allometry often has the strongest effect on shape variation (Bright et al., 2016). Similar results have been found in broad taxonomic studies, with as little as 12% of variation in beak shape associated with diet (Navalon et al., 2019).

In light of these contradictory findings, it has been suggested that traditional dietary categories are too broad to capture the diversity of function in bird skulls (Pigot et al., 2016; Navalon et al., 2019; Felice et al., 2019). In a study mapping avian morphology to associated trophic niche across the breadth of extant bird diversity, a minimum of four morphological trait dimensions were required to parse out phylogenetic noise or convergence of form (Pigot et al., 2020). Likewise, a study successfully linking skull shape with foraging ecology in Charadriiformes (shorebirds, gulls, and auks) found that after collapsing the number of foraging guilds from 36 to 10, the explanatory power of foraging ecology decreased by nearly 50% (Natale & Slater, 2022). Together, these results suggest the need for more descriptive categorization linking feeding ecology and form.

Vultures are a paraphyletic functional guild formed from members of two avian families; Afro-Eurasian vultures (Accipitridae), and American vultures (Cathartidae; Jarvis et al., 2014). Obligate scavenging in vertebrates is only found in vultures, and it has evolved independently in these two families. Convergent evolution appears to have favored a number of highly specialized traits adapted to foraging for carrion. These birds share exceptionally keen eyesight, specialized digestive tracts, and soaring flight, allowing them to easily locate and rapidly consume detritus material (Ruxton & Houston, 2004; Potier, 2020; Ogada et al., 2012; Kane & Kendall, 2017; Houston, 1975). Given the high apparent convergence of attributes in this specialized ecological guild, a strong link between physical form and dietary preference might be expected. Vultures thus represent an interesting model for investigating the degree to which ecological and evolutionary factors contribute to variation in skull shape.

Across the 23 extant species, this guild exhibits phylogenetic (Jarvis et al., 2014), ecological (van Overveld et al., 2020; Linde-Medina et al., 2021), and morphological diversity (Hertel, 1994; Böhmer et al., 2020; Holmes et al., 2022). Distinctions in sociality (Kendall, 2013; van Overveld et al., 2020), breeding and nesting behavior (Kemp & Kemp, 1975; Krüger et al., 2015; Kendall, 2013; Mundy et al., 1992), migratory and movement patterns (Alarcón & Lambertucci, 2018), habitat preferences (Kendall, 2014; Del Hoyo et al., 1992), sensory perception (Spiegel et al., 2013; Portugal et al., 2017; Ogada et al., 2012; Jackson et al., 2020), and feeding and foraging strategies (Kruuk, 1967; Houston, 1987; Ogada et al., 2012; Jackson et al., 2020; van Overveld et al., 2022) have been recorded. For example, *Gypohierax angolensis* and *Gypaetus barbatus* both display unique dietary preferences, with the former primarily an herbivore (Lambertucci et al., 2021) and the latter a bone specialist (Cramp, 1980). Intense competition for spatially and temporally unpredictable food has likely driven many of these differences (Böhmer et al., 2020; Holland et al., 2019). Diversity among vultures is a strong base for testing competing hypotheses for underlying drivers of avian cranial morphology.

Previous behavioral research has provided evidence that vultures fall into three distinct ecotypes based on mode of feeding and dietary preference; ripper, gulper, and scrapper (Kruuk, 1967; König, 1974; König, 1983; Houston, 1987; Hertel, 1994; Table 1). Additionally, there is evidence that these ecotypes are reflected in the anatomy of the skull (Hertel, 1994) and neck (Böhmer et al., 2020). Morphometric analyses investigating the effects of diet on raptor skull morphology have consistently placed vultures outside other groups, even in the absence of other dietary trends and to the point of occupying an almost entirely isolated region of morphospace (Hertel, 1995; Guangdi et al., 2015; Bright et al., 2016; Sun et al., 2018; Pecsics et al., 2019). These studies typically classify vultures as “scavengers,” preventing distinctions being made on the basis of various feeding strategies. Few studies have investigated skull shape variation across vulture feeding types specifically (Hertel, 1994; Linde-Medina et al., 2021), and those that have relied on traditional methods of linear measurements, which omit detailed shape information (Goswami et al., 2019). Geometric morphometric methods allow the accurate quantification of shape, outperforming traditional methods in both accuracy and detail (Maderbacher et al., 2008; Breno et al., 2011; Mendonca et al., 2013; Parés-Casanova et al., 2020), and allowing for the visualization of shape variation (Breno et al., 2011; Parés-Casanova et al., 2020).

**Table 1.** Vulture Feeding Classification System.

| Feeding Type | Diet | Mode of Feeding |
| --- | --- | --- |
| **Ripper** | Tougher material, skin/hide, muscle, and tendons. Typically feed on the external areas of a carcass. | Strong tearing action away from the carcass. |
| **Gulper** | Soft tissue, and viscera. Typically feed on the internal material of a carcass. | Complete insertion of the head into the carcass for swallowing soft food. |
| **Scraper** | Scraps of meat found around the carcass, often the leftover material of another feeding scavenger. | Pecking motion to pick up small scraps on the ground and around the carcass. |

Here, we investigate the relative contributions of allometry, phylogeny, and vulture feeding type on variation in skull shape using three-dimensional geometric morphometrics. We predict that vulture skull shape is correlated with feeding ecology. We expect allometry to have a greater influence on skull shape than feeding ecology, because there is evidence to suggest that skull shape in raptors is highly integrated with size as an adaptive strategy for rapid evolution (Bright et al., 2016). Finally, because of convergence in feeding ecologies across family groups, we expect that the phylogenetic signal in skull shape is low, and phenotypic convergence is high (Jarvis et al., 2014; Linde- Medina et al., 2021).

## Materials and Methods

We quantified skull morphology in a dataset composed of 22 extant vulture species, one extinct vulture, and eight non-vulture raptors (Supplementary Table S1). Non-vulture raptors were selected on the basis of sharing similarities in body size, ecology (generalists and frequent scavengers), and geographic overlap in range (Cramp, 1980; Blem, 1997). Three-dimensional meshes were created from a total of 31 specimens (one representative per species) obtained from MorphoSource (www.morphosource.org), Phenome 10K (www.phenome10k.org), Sketchfab (www.sketchfab.com), personal communications, or directly scanned from museum collections (Supplementary Table S2). All specimens were analyzed without the rhamphotheca, a layer of keratin that covers the beak, because this is part of the epidermis and not osseous skull material. No mandibles were included in this study.

Meshes were processed with Geomagic Wrap 2017 (3D Systems Inc., Rock Hill, SC, USA) to remove scanning artifacts and fill holes. Each mesh was landmarked with 38 anatomical landmarks and 24 sliding semi-landmark curves in Stratovan Checkpoint (Stratovan Corporation, Davis, CA, USA), using a template adapted from Mitchell et al. (2021; Supplementary Tables S3 and S4). Anatomical landmarks were placed bilaterally and semi-landmark curves were placed on the right side. Damaged specimens were mirrored in Geomagic Wrap before landmarking. Semi-landmark curves were slid to minimize bending energy (Gunz et al., 2005) with an adaptation of the “slider3d” function in the *morpho* package in R (Schlager, 2017; R Core Team, 2021.). Right-side landmarks were temporarily mirrored to the left side of the specimen during Procrustes alignment to avoid introducing error and to improve estimates of shape variation and allometry (Cardini, 2016). Mirroring was done with the “mirrorfill” function from the *paleomorph* R package (Lucas & Goswami, 2017). Landmark data were then superimposed with a generalized Procrustes alignment (GPA) (Rohlf & Slice, 1990) to minimize differences in size, orientation, and location between landmark sets (Kendall, 1989) with the “gpagen” function in *geomorph* (Baken et al., 2021; Adams et al., 2022). Analyses were repeated on separately aligned subsets of the data to account for non-vultures and unknown ecological categories (See supplementary materials). Left-side landmarks were removed after alignment, leaving a total of 359 landmarks per specimen. A principal components analysis (PCA) was performed on the Procrustes-aligned shape data to explore shape variation (Collyer & Adams, 2021).

We generated a time-scaled phylogeny from BirdTree.org (Jetz et al., 2012) based on Hackett et al. (2008; Hackett All Species: a set of 10000 trees with 9993 OTUs each) for all birds of prey (Accipitridae, Pandionidae, Sagittariidae, Falconidae, Cathartidae, and Cariamidae). The resulting tree was pruned in Mesquite (Madison & Madison, 2021) to our dataset. The extinct *Hieraaetus moorei* was substituted in the place of its closest living relative, *Hieraaetus morphnoides* (Bunce et al., 2005). The extinct *Breagyps clarki* was added as a sister taxon to *Gymnogyps californianus* (Emslie, 1988), with the node placed midway along the branch subtending *G. californianus*. The resulting time-scaled tree was read into R using the *ape* package (Paradis & Schliep, 2019). We calculated phylogenetic signal (the degree of similarity explained by shared ancestry) using the *K_mult_* statistic, implemented with the “physignal” function in *geomorph* (Adams, 2014a). Allometric influence on skull shape was tested with raw shape data and phylogenetically corrected shape data respectively with the “procD.lm” and “procD.pgls” functions in *geomorph* (Anderson, 2001; Adams, 2014b).

Feeding type was assigned to each species following the classification scheme created by Hertel (1994; Table 1) and based on behavioral observations in the field (Kruuk, 1967; König, 1974; König, 1983; Houston, 1987; Hille et al., 2016; Gaengler & Clum, 2015; J. Burnett, pers. comms.) with the exception of the *Gypohierax angolensis* and *Gypaetus barbatus*, which do not fit these categories (König, 1974; Linde-Medina et al., 2021). These two species along with all extinct and non-vulture raptors were not assigned a type (Figure 1). A multivariate analysis of variance (MANOVA) was performed on the Procrustes-aligned shape data with feeding type as the independent grouping variable to determine if skull shape correlated with feeding groups. This was repeated for allometry-corrected shape values to identify a potentially significant interaction with size. Interactions between allometry and feeding type as well as phylogeny and feeding type were explored with the “procD.lm” (for raw shape data) and “procD.pgls” (for phylogenetically corrected shape data) functions in *geomorph* (Anderson, 2001; Adams, 2014b). Feeding types were plotted over principal component scores in the morphospace. To test the fit of the shape data with feeding categories, we implemented a discriminant function analysis (DFA) with the ‘mvgls.dfa’ function in the R package *mvMORPH* (Clavel et al., 2015). This method computes a discriminant analysis based on GLS estimates from a phylogenetic regression model and is optimized for high–dimensional data. The output shows both assignment accuracy of specimens of known feeding groups and predicts group assignment for specimens of unassigned group, including the two extant vulture species that did not fall into any of the three feeding categories (*G. angolensis* and *G. barbatus*), and the extinct vulture, *B. clarki*. Phenotypic convergence within feeding groups was quantified with the distance-based measure, C, developed by Stayton (2015), an approach which measures the average phenotypic convergence across a group within phylomorphospace. Results were compared with a set of 100 simulations under a BM null model of evolution to provide a significance value for each cluster, using the first 14 principal components, accounting for a cumulative 95% of total shape disparity, and implemented with the ‘convratsig’ function in the R package *convevol* (v. 2.0.0, Stayton, 2015). This method has been shown to overestimate convergence in outlying taxa, and so we also applied the method developed by Grossnickle et al. (2023), Ct, which measures phenotypic distance at equivalent points on a time-scaled phylogeny. This was implemented in the R package *convevol* (Stayton, 2015), using the same parameters as the C measures calculated above.

**Figure 1:**
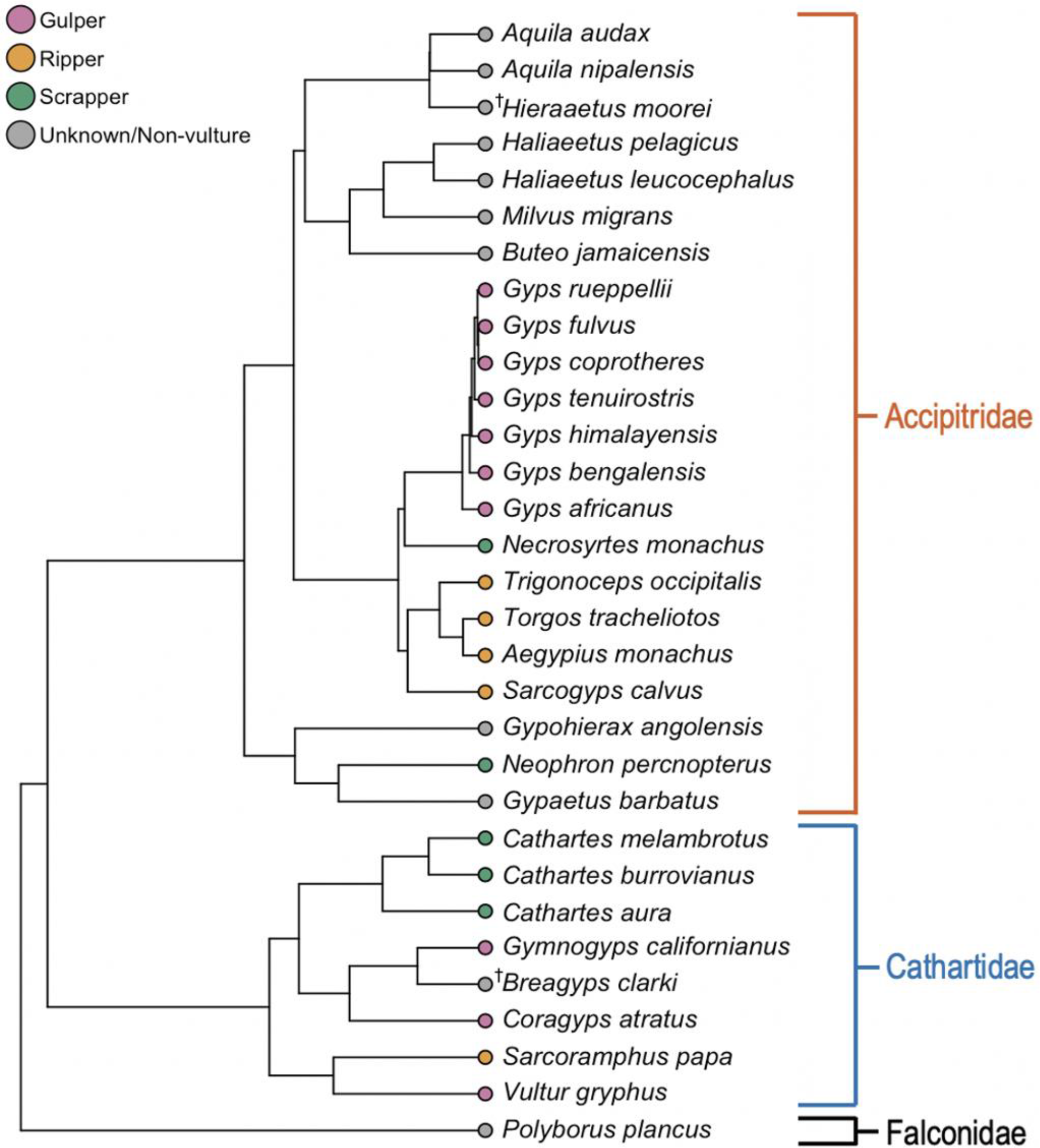
Phylogeny of the 31 species included in this study, adapted from BirdTree.org (www.birdtree.org) (Jetz et al., 2012) based on Hackett et al. (2008). Taxa are colored according to feeding type. Extinct species are marked (e.g. ^†^*Breagyps clarki*).

**Figure 2.**
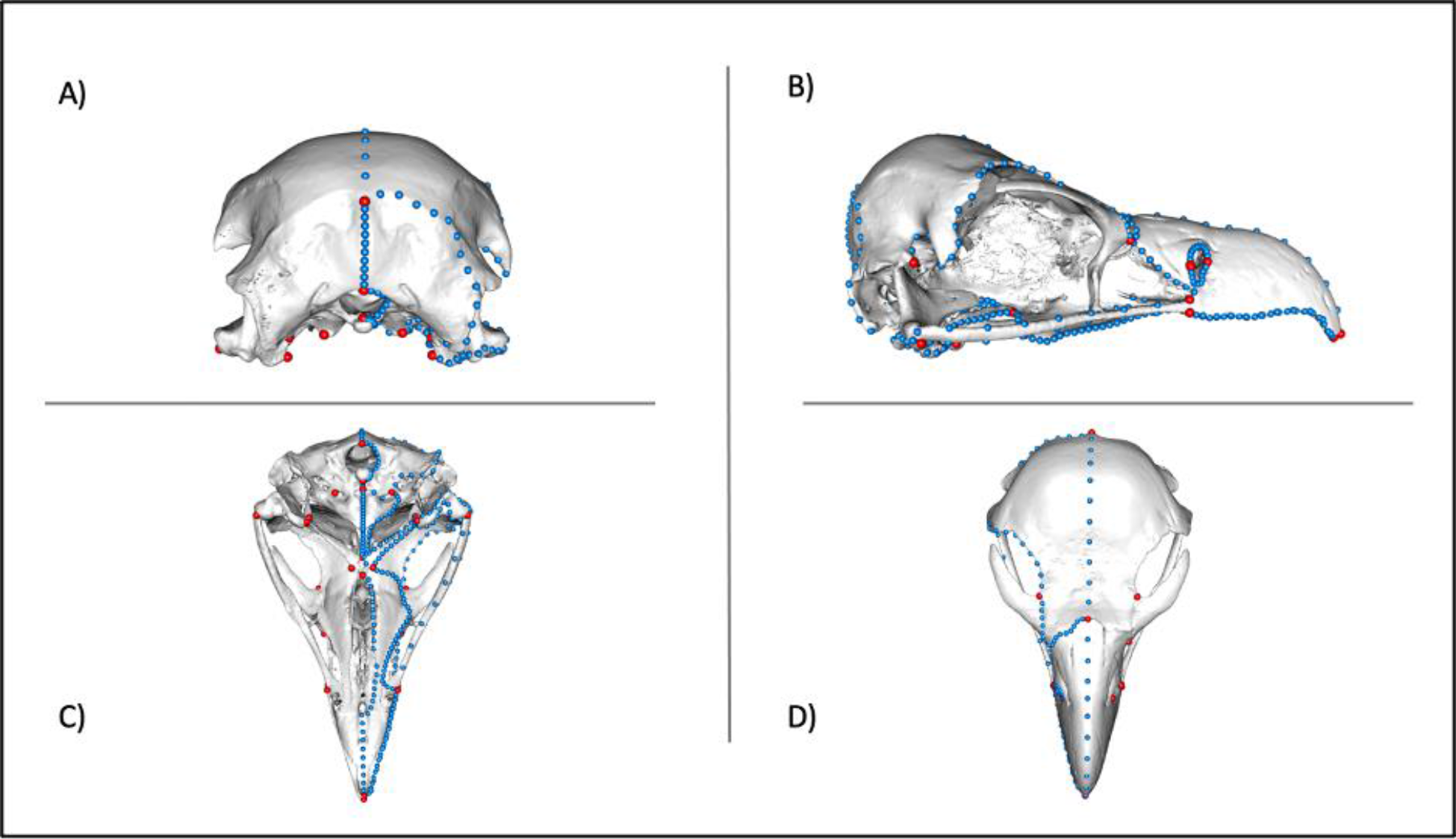
Configuration of the 38 anatomical landmarks (red) and 24 semi-landmark curves (blue) used in this study. Landmarks are visualized on a three-dimensional mesh of *Trigonoceps occipitalis* with A) posterior, B) lateral, C) ventral, and D) dorsal views shown.

## Results

In the full dataset, the first 14 principal components (PCs) account for ∼95% of total shape variation, with the first three PCs cumulatively accounting for ∼65% of shape variation (Supplementary Figure S1). Shape change along PC1 (proportional variance: 35.8%) is characterized by a transition from a short to elongate beak and naris, tall to short cranium, and an increasingly laterally orientated orbit. Shape change along PC2 (proportional variance: 19.8%) is characterized by a transition from slender to robust beak, an increasingly angular craniofacial hinge, and a large, elongate oval naris to a thin, vertical naris opening. Accipitridae and Cathartidae separate out along PC2. Accipitrids are generally characterized by a comparatively taller and wider cranium, shorter and slimmer nares, a more angular craniofacial hinge, and robust beaks. Cathartids are characterized by a lower skull, elongate and slender beaks, longer nares, and a distinctly anteriorly sloping cranium. The width of the frontal bone tends to be thinner in accipitrid species anterior to the postorbital process, at which point it expands more laterally than in cathartid taxa. Shape change along PC3 (proportional variance: 9.6%) occurs almost entirely in the beak by shifting towards a more robust and deeply hooked beak (Figure 3). Phylogenetic signal in shape data was moderately low but statistically significant (*K_mult_* = 0.325, *p* = 0.001, Supplementary Table S6), suggesting a degree of convergence in shape within the dataset. Allometry accounts for 18.5% of total shape variation (*R^2^* = 0.185, *Z* = 3.631, *p* = 0.001; Supplementary Table S7), but is not significant after accounting for phylogeny (*R^2^* = 0.05, *Z* = 1.463, *p* = 0.073; Supplementary Table S7).

**Figure 3.**
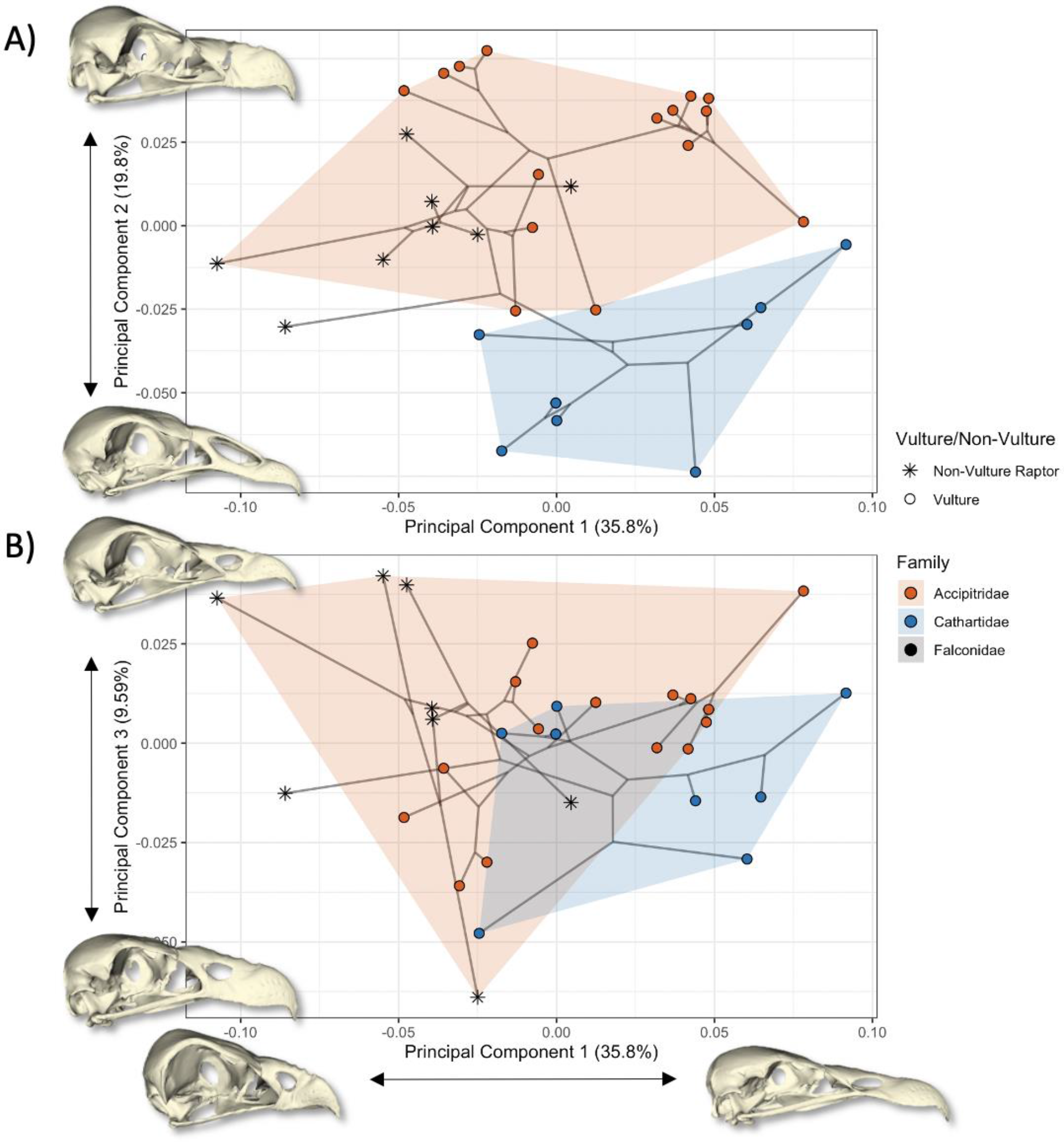
Phylomorphospace showing avian families overlaying the species in this study with transformation along PC1 on the x axis and transformations along PC2 (A) and PC3 (B) on the y axes. Warped meshes show positive and negative shape values along each axis.

Feeding groups mapped over PCs 1 and 2 occupy distinct regions of morphospace, with no overlap along PC1 (Figure 4). Feeding type is significantly correlated with skull shape in non-phylogenetically corrected shape data (*R^2^* = 0.457, *Z* = 4.789, *p* = 0.001; Supplementary Table S7), but this correlation disappears after correcting for phylogenetic relatedness (*R^2^* = 0.089, *Z* = 0.159, *p* = 0.436; Supplementary Table S7). Mean phenotypic convergence is statistically significant within all feeding groups with the *C* measure of phenotypic convergence (Stayton, 2015; gulper *C_1_* = 0.39, *p* = <0.001; ripper *C_1_* = 0.33, *p* = <0.001; scrapper *C_1_* = 0.35, *p* = <0.001). Convergence is not significant across any group when implementing the Ct measure (Grossnickle et al., 2023; ripper, *Ct_1_* = -0.296, *p* = 0.47; gulper, *Ct_1_* = -0.978, *p* = 0.55; scrapper, *Ct_1_* = -0.105, *p* = 0.05).

**Figure 4.**
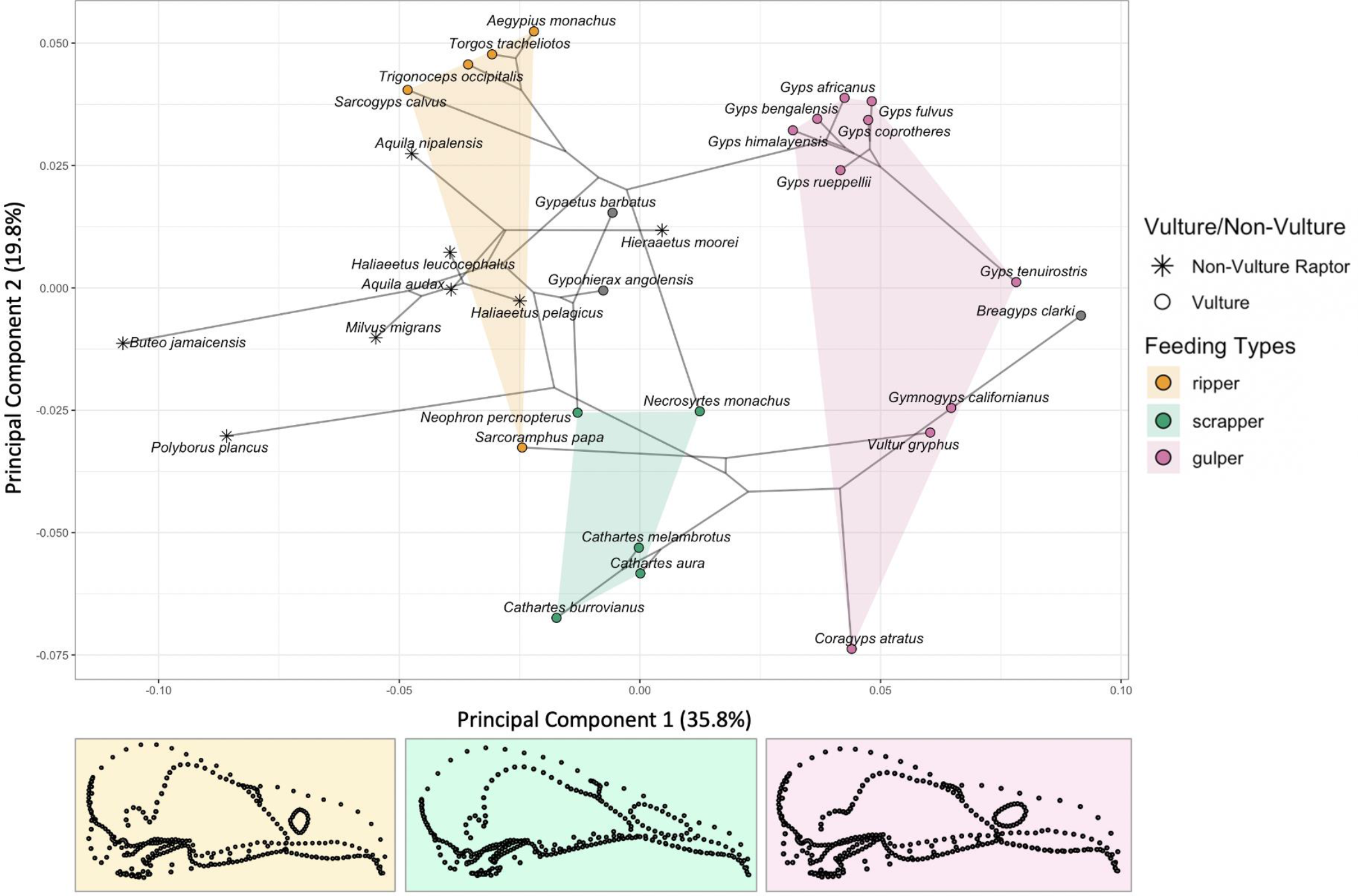
Phylomorphospace of PCs 1 and 2 with vulture feeding types grouped by color and non-vultures and extinct *Breagyps clarki* shown in gray. The color-coded mean shape for each feeding type is shown below the figure.

When examining putatively converging taxa rather than all taxa within feeding groups, however, convergence was significant and high for gulper (*Vultur gryphus, Gymnogyps californianus, Gyps tenuirostris*; *Ct_1_* = 0.63, *p* = 0.00, Supplementary Figure S2) and scrapper (*Necrosyrtes monachus, Neophron percnopterus, Cathartes melambratus*; *Ct_1_* = 0.45, *p* = 0.00, Supplementary Figure S3) taxa, but not for ripper taxa (*Sarcoramphus papa, Torgos tracheliotos*; *Ct_1_* = -0.004, *p* = 0.38, Supplementary Figure S4).

The DFA assigned 100% of specimens of known feeding ecology to the correct group (Supplementary Table S8). The two unassigned extant species (*Gypohierax angolensis* and *Gypaetus barbatus*) were assigned to the scrapper group (Supplementary Table S8). The extinct *Breagyps clarki* was assigned to the gulper group. Upon including a fourth feeding group for non-vulture raptors, the predicted group assignment for *G. barbatus* changed from scrapper to non-vulture raptor (Supplementary Table S9).

## Discussion

Vulture skull shape is influenced by the interplay of a variety of factors, mainly phylogeny, feeding strategy, and allometry. The distinct shape of vulture skulls compared to non- vulture raptors, coupled with examples of convergence in shape between vultures from both families, suggest that the unique feeding ecologies of vultures are considerable drivers of vulture skull evolution, and highlight the evolutionary constraints of ecologically specialized taxa (Bennett & Owens, 1997). Although vulture skull shape is not significantly correlated with feeding ecology after accounting for phylogeny, feeding groups occupy distinct regions of morphospace (Figure 4), suggesting that adaptation to different feeding ecologies has driven divergence in skull shape within Accipitridae and Cathartidae. This is supported by examples of phenotypic convergence and parallel evolution between these families, and skull shape consequently performs very well in predicting feeding ecology. These results reinforce popular hypotheses that the evolution of bird skulls is driven by dietary needs (Lack, 1953; Gibbs & Grant, 1987; Lovette et al., 2002; Jønsson et al., 2012), as well as support recent evidence in the literature that incorporating finer detail within a smaller phylogenetic context could provide more information on the relationship between form and function (Pigot et al., 2016; Olsen, 2017; Navalón et al., 2019; Felice et al., 2019; Pigot et al., 2020; Natale & Slater, 2022).

Skull shape variation across the data set yielded a significant, albeit moderately low, phylogenetic signal (*K_mult_* = 0.325), revealing that shape is phylogenetically structured, but that phenotypic convergence and parallel evolution play an important role in vulture skull evolution. The separation of Accipitridae and Cathartidae along PC2 reveals the distinct morphologies of each family. Most notably, accipitrid skulls tend to be tall and robust while cathartid skulls are low and slender. The description “low and slender” is commonly used to differentiate all vultures, accipitrids included, from non-scavenging raptors (Hertel, 1994; Guangdi et al., 2015; Pecsics et al., 2019), underscoring the importance of describing feeding ecology in finer detail. Some distinguishing features of the accipitrid skull could offer advantages in the predominantly open, grassland habitats of these vultures (Holmes et al., 2022). Accipitrid vultures tend to soar at higher altitudes and consume larger carcasses than cathartids (Mundy et al., 1992; Houston, 1984), relying primarily on vision to locate both conspecifics and carcasses (Dermody et al., 2011; Potier, 2020). The comparatively larger orbits of accipitrid skulls may indicate greater visual acuity (Potier, 2020; Ogada et al., 2012). Another distinction of the accipitrid skull is a smaller naris, a feature particularly striking in *Gyps* species whose nostrils are partly covered by a bony sheath, leaving a thin vertical opening. No explanation for these sheaths exists in the literature, though protection from dust in semi-arid habitats, viscera when feeding, or wind at high altitudes are all possible explanations. Conversely, visual abilities may be of limited use to American vultures in their often densely forested habitats, thus a reliance on olfaction to locate food has likely driven the large, open nares of American vultures (Ogada et al., 2012; Houston, 1984; Houston, 1987).

In raw shape data, feeding group explained the highest proportion (∼45%) of shape disparity, seemingly providing strong evidence that the evolution of skull shape in vultures is driven by feeding behavior. As hypothesized by Hertel (1994), vultures fall into three distinct regions of morphospace based on feeding strategy, with no overlap along PC1 (Figure 4). When phylogenetic relatedness is accounted for, however, feeding type is not significantly correlated with shape (*p* = 0.436). This is probably due to clusters of closely related species that share feeding ecologies (e.g. *Gyps*, seven species which are all gulpers) overwhelming the convergence signal of smaller numbers of more distantly-related taxa. Nonetheless, there are clear examples of ecological (Figure 1) and phenotypic convergence (Figure 4) in the dataset. Notably, *Neophron percnopterus* and *Necrosyrtes monachus*, two accipitrids, converge on Cathartidae taxa along PC2 in the ‘scrapper’ region of morphospace. For both species, the most closely related taxa do not share the same feeding ecology. Similarly, *Sarcoramphus papa* and *Vultur gryphus* fall into ‘ripper’ and ‘gulper’ space respectively, demonstrating strong morphological divergence in these sister taxa. Using the C measure of Stayton (2015), within-group phenotypic convergence was found to be significant, and relatively consistent within all feeding groups, with an average of 39% convergence in gulpers, 33% in rippers and 35% in scrappers. This is contradicted by the results of the Ct measure of phenotypic convergence of Grossnickle et al. (2023), with no groups showing significant convergence overall. Examples of significantly converging taxa, however, can be found by focusing on smaller numbers of taxa that appear to show convergence within morphospace, rather than across whole groups. The most conspicuous example of convergence found with this method occurs between *V. gryphus*, *Gymnogyps californianus*, and *Gyps tenuirostris* which, despite large phylogenetic distance, have converged on the same region of morphospace (*Ct_1_* = 0.63, *p* = 0.00; Figure 4). The ripper feeding group did not show any significant convergence with the Ct method, despite being clearly separated from other feeding groups along PC1. This is likely to be a combination of the majority of taxa in this group being closely related accipitrids and thus more divergent, with only one more distantly related cathartid species in this group, *S. papa*, ‘converging’ on this region of morphospace. The Ct method only measures phenotypic distance at equivalent timesteps on a time-scaled phylogeny, and so examples of parallel evolution, which can superficially resemble convergent evolution in some circumstances, are not recognised as convergent with this method. This contrasts with the C measure, which often cannot differentiate between convergent and parallel evolution (Grossnickle et al., 2023). Nonetheless, the apparent parallel evolution along PC1 of *S. papa* with the ripper taxa in Accipitridae is notable in that the evolution of this feeding type coincides with a marked negative shift along PC1.

The mean shapes generated from these groups (Figure 4) reflect what might be expected by each vulture feeding type. ‘Rippers’ have a wider cranium and more robust beak for tearing tougher tissue from carcasses. ‘Gulpers’ have the narrowest skull with the relatively longest beak, supporting ease of maneuverability inside a carcass. ‘Scrappers’ have the slenderest beak, reflecting the precision necessary for picking up small scraps around the carcass. In most other respects, the ‘scrapper’ shape is intermediate to the other types, in accordance with the more generalist strategies of various scrapper species (Ballejo et al., 2018).

Although large bodies allow vultures to maximize food consumption at spatially and temporally unpredictable food sources, the mechanism selecting for and constraining this ability is soaring flight (Ruxton & Houston, 2004; Poessel et al., 2018), rather than feeding behavior. This likely explains the lack of correlation between skull size and shape among vultures. Thus, body size in vultures probably evolved in response to selective pressures acting on searching or foraging efficiency such as flight conditions (Ruxton & Houston, 2004; Houston 1987), habitat (Xirouchakis & Mylonas, 2004), species interactions (van Overveld et al., 2020; van Overveld et al., 2022; Jackson et al., 2020), and physiological capacity (Ruxton & Houston, 2004). Future morphometric research investigating the relationship between vulture feeding types and other ecological traits, particularly species interactions, is recommended.

A handful of species included in the study either do not have sufficient behavioral observations to support a vulture feeding type assignment or have been contested in the literature. *Coragyps atratus*, originally identified as a ‘gulper’ based on behavioral observations (Houston, 1987), has been predicted by morphological research as both a ‘scrapper’ (Hertel, 1994) and a ‘gulper’ (Linde-Medina et al., 2021). The overlap with ‘gulpers’ along PC1 and ‘scrappers’ along PC2 highlights the limitations of studies based on morphology alone, and the importance of supplementing morphometric data with behavioral observations. Based on morphology alone, *Gypaetus barbatus* has previously been classified as a gulper (Hertel, 1994) and ripper (Linde-Medina et al., 2021), and *Gypohierax angolensis* has been proposed as both a gulper (Hertel, 1994) and scrapper (Linde-Medina et al., 2021). Using a discriminant function analysis to predict unknowns, as both Hertel (1994) and Linde-Medina et al. (2021) did, our study classified both species as scrappers (Supplementary Table S8). When including non-vulture raptors in the model, however, *G. barbatus* was reassigned as a raptor (Supplementary Table S9). These discrepancies further highlight the limitations of morphology-based predictions and the risks of overriding observed behavior. The intermediate positions of these species in morphospace as well as their proximity to non-vulture raptors, suggest that these two species have not undergone such extreme morphological evolution as other, more specialized vultures. In addition, although the ‘gulper’ assignment of *Gyps tenuirostris* is supported by field observations (Hille et al., 2016), this is the first morphometric study on vulture feeding types to include this taxon and confirm morphological similarities with other ‘gulpers,’ including the distantly related cathartids *Vultur gryphus* and *Gymnogyps californianus* (Figure 4). Finally, it is possible to extrapolate feeding assignments to extinct species using discriminant analyses, though results should be interpreted with caution given the inability to obtain observational feeding data. The results of our DFA matched those of Hertel (1994), predicting *Breagyps clarki* a gulper regardless of the inclusion of a raptor category (Supplementary Tables S8 and S9). The extinct *Hieraetus moorei* has been the subject of debate regarding its feeding ecology and was recently proposed a ‘gulper’ on account of its morphological similarity to *V. gryphus* (Van Heteren et al., 2021). In contradiction with the obligate scavenger hypothesis, our study finds no morphological evidence to support a vulture feeding type assignment for this species, and based on hindlimb morphology was almost certainly a raptor (Van Heteren et al., 2021). Our DFA predicted this species to be a ‘scrapper’, but was assigned as a ‘raptor’ when non–vulture raptors were included (Supplementary Tables S8 and S9). A better understanding of non-vulture raptor feeding ecology will improve feeding classification and prediction within this group.

The ability to predict function from form has been a contentious topic in bird skull morphometrics (Navalón et al., 2019; Natale & Slater, 2022), and to do so using a single functional trait has had mixed results (Pigot et al., 2020; Ballentine et al., 2013). The combined dietary and biomechanical information encoded in vulture feeding types is one possible explanation for drawing a successful link between feeding ecology and skull shape. Providing more detail of ecological context has been shown to improve predictive power in morphometric studies (Pigot et al., 2016; Navalón et al., 2019; Friedman et al., 2019) although the task of handling one-to-many or many-to-one ecomorphological relationships remains a challenge (Pigot et al., 2016; Navalón et al., 2019; Friedman et al., 2019). The focus on a single functional guild improved the detail and accuracy of functional traits, supporting the idea that taxonomic categories are possibly too broad to provide meaningful results in such an ecologically and phenotypically diverse class (Felice et al., 2019; Pigot et al., 2020). “Scavenger” is a broad term applicable to many opportunist and carnivorous species (DeVault et al., 2003), and inclusion under this umbrella term has the potential to group together specialized (ecologically constrained) and generalist (ecologically flexible) taxa.

## Conclusion

The use of geometric morphometrics to investigate the evolution and diversification of the avian cranium has yielded new and unexpected discoveries into the various factors contributing to shape variation, while casting doubt on traditional associations between beak shape and ecological niche (Felice et al., 2019; Bright et al., 2016; Navalón et al., 2019; Bright et al., 2019; Tattersall et al., 2017). The avian beak is a multi-functional apparatus, however, and a complex variety of selective pressures influence the tempo, direction, and mode of avian skull morphology, both developmentally and ecologically (Felice et al., 2019). Broadly applying hypotheses across Aves is likely to provide equally complex results. The ability to link feeding ecology, rather than broader dietary categories to skull shape in the present study is a potentially fruitful avenue of research. Future research testing further competing hypotheses on vulture skull shape variation in relation to cross-species interactions and functional traits is recommended, as this could offer additional insights into the evolution of obligate scavenging. The results of this study also have important implications for the conservation of this rapidly declining guild (Ogada et al., 2012) as vulture conservation initiatives often involve the use of artificially provided food sources such as supplementary feeding sites (Margalida et al., 2010). Furthermore, future research on internal skull morphology could highlight key differences in sensory perception, both at the species level and between feeding types, which would allow for more reliable predictions for human-induced change (Martin et al., 2012).

## Acknowledgements

The authors thank Joe Burnett (Ventana Wildlife Society), Daniel Field (University of Cambridge), Oliver Kippax-Chui (Natural History Museum), Ryan Marek (University College London), Paul Scofield (University of Canterbury), and Judith White (Natural History Museum) for helping with data collection. The authors thank the Goswami Lab at the Natural History Museum in London for useful discussions and help with processing and analyzing data.

## Conflict of Interest

The authors declare no conflict of interest.

## Author Contributions

KRS and AK conceived the study; KRS, MKS and AK collected the data; All authors performed the analyses and wrote the manuscript.

## Notes

### Competing Interest Statement

The authors have declared no competing interest.

