## Supplementary tables and figures for "Carrion converging: Skull shape is predicted by feeding ecology in vultures"

**Supplementary Materials.**

**Katherine R Steinfield, Ryan N Felice, Mackenzie E Kirchner and Andrew Knapp**

**Table S1. List of study species.** Study species included in analyses alongside taxonomy, body mass (Dunning, 2007), and feeding type assignments as per Hertel (1994) supported by field observations (Kruuk, 1967; König, 1974; König, 1983; Houston, 1987; Hille et al., 2016; Gaengler & Clum, 2015; Burnett, pers. comms.). Institutional codes correspond to collections at The Natural History Museum, London (NHMUK), the University Museum of Zoology, University of Cambridge (UMZ), the Smithsonian National Museum of Natural History (USNM), the University of California Museum of Vertebrate Zoology (UCMVZ), the University of California Museum of Paleontology (UCMP), the Canterbury Museum (CMC), the Muséum National d’Histoire Naturelle (MNHN), and the Carnegie Museum of Natural History (CMNH).

| Species | Specimen number | Family | Mass (g) | Feeding Group |
| --- | --- | --- | --- | --- |
| *Aegypius monachus* | NHMUK 1872.10.25.5 | Accipitridae | 9625 | ripper |
| *Aquila audax* | NHMUK S.1966.51.16 | Accipitridae | 3466 | raptor |
| *Aquila nipalensis* | NHMUK 1923.9.3.1 | Accipitridae | 2746 | raptor |
| *Breagyps clarki* | UCMP 299082 & 241455* | Cathartidae | NA | NA |
| *Buteo jamaicensis* | UCMVZ 180162 | Accipitridae | 1126 | raptor |
| *Cathartes aura* | USNM 17872 | Cathartidae | 2006 | scrapper |
| *Cathartes burrovianus* | USNM 622341 | Cathartidae | 935 | scrapper |
| *Cathartes melambrotus* | USNM 621939 | Cathartidae | 1373 | scrapper |
| *Coragyps atratus* | UCMVZ 78681 | Cathartidae | 2159 | scrapper |
| *Gymnogyps californianus* | UCMVZ 151087 | Cathartidae | 8450 | gulper |
| *Gypaetus barbatus* | NHMUK S.1962.7.1 | Accipitridae | 5680 | NA |
| *Gypohierax angolensis* | NHMUK 224820 | Accipitridae | 1600 | NA |
| *Gyps africanus* | USNM 430014 | Accipitridae | 5433 | gulper |
| *Gyps bengalensis* | NHMUK 2004.2.3 | Accipitridae | 4385 | gulper |
| *Gyps coprotheres* | USNM 561314 | Accipitridae | 8177 | gulper |
| *Gyps fulvus* | MNHN 1995-160 | Accipitridae | 7436 | gulper |
| *Gyps himalayensis* | USNM 19534 | Accipitridae | 10000 | gulper |
| *Gyps rueppellii* | USNM 430178 | Accipitridae | 7400 | gulper |
| *Gyps tenuirostris*** | UMZ 13.Acc.25.d.2 | Accipitridae | 5515 | gulper |
| *Haliaeetus leucocephalus* | NHMUK 1869.10.19.2 | Accipitridae | 4740 | raptor |
| *Haliaeetus pelagicus* | NHMUK S.1996.31.1 | Accipitridae | 7757 | raptor |
| *Hieraaetus moorei* | CMC AV5684 | Accipitridae | NA | NA |
| *Milvus migrans* | UCMVZ 115927 | Accipitridae | 567 | raptor |
| *Necrosyrtes monachus**** | CMNH A6841 | Accipitridae | 2043 | scrapper |
| *Neophron percnopterus* | USNM 17835 | Accipitridae | 2082 | scrapper |
| *Polyborus plancus* | USNM 227375 | Falconidae | 1348 | raptor |
| *Sarcogyps calvus* | NHMUK S.2013.22.1 | Accipitridae | 4550 | ripper |
| *Sarcoramphus papa* | MNHN A4002-IV/188 | Cathartidae | 3400 | ripper |
| *Torgos tracheliotos* | NHMUK S.1952.1.172 | Accipitridae | 6969 | ripper |
| *Trigonoceps occipitalis* | USNM 320859.2 | Accipitridae | 3016 | ripper |
| *Vultur gryphus* | NHMUK S.1956.1.81 | Cathartidae | 11300 | gulper |

*The two specimen IDs correspond to two parts of the same individual.

**This specimen is identified in UMZ collections as *Gyps indicus*. However, Mundy (2022) re-assigned this specimen as *Gyps tenuirostris*.

***Available for restricted download from Morphosource (<https://www.morphosource.org>, media ID: 113601).

**Table S3. Anatomical landmark guide.** Adapted from Mitchell et al., 2021.

| Landmark | Description |
| --- | --- |
| 1 | Tip of rostrum, palatal surface, midline |
| 2 | Posterior most point of choana at midline |
| 3 | Ventral contact between premaxilla and jugal bar (right side) |
| 4 | Tip of rostrum, anterodorsal side, midline |
| 5 | Median point of craniofacial hinge |
| 6 | Anterodorsal limit of the frontal contribution to the orbit-lacrimal contact (right side) |
| 7 | Anteroventral contact between pterygoid and palatal surface (right side) |
| 8 | Tip of zygomatic process of squamosal (right side) |
| 9 | Medial point of contact between parietal and supraoccipital |
| 10 | Medial point of dorsal margin of foramen magnum |
| 11 | Medial point of dorsal surface occipital condyle |
| 12 | Medial point of ventral surface occipital condyle |
| 13 | Medial point of contact between basicoccipital and basisphenoid |
| 14 | Anterior most point of basisphenoid, just posterior to the pterygoids and palate |
| 15 | Posteromedial corner of articular process of quadrate (right side) |
| 16 | Anterolateral corner of articular process of quadrate (right side) |
| 17 | Posterior point of pterygoid-quadrate articulation (right side) |
| 18 | Anterior point of pterygoid-quadrate articulation (right side) |
| 19 | Anterior most point of external naris (right side) |
| 20 | Posterior most point of ventral (lateral) margin of external naris (right side) |
| 21 | Posterior most point of dorsal (medial) margin of external naris (right side) |
| 22 | Lateral extreme of frontonasal contact (right side) |
| 23 | Ventral contact between premaxilla and jugal bar (left side) |
| 24 | Anterodorsal limit of the frontal contribution to the orbit-lacrimal contact (left side) |
| 25 | Anteroventral contact between pterygoid and palatal surface (left side) |
| 26 | Tip of zygomatic process of squamosal (left side) |
| 27 | Posteromedial corner of articular process of quadrate (left side) |
| 28 | Anterolateral corner of articular process of quadrate (left side) |
| 29 | Posterior point of pterygoid-quadrate articulation (left side) |
| 30 | Anterior point of pterygoid-quadrate articulation (left side) |
| 31 | Anterior most point of external naris (left side) |
| 32 | Posterior most point of ventral (lateral) margin of external naris (left side) |
| 33 | Posterior most point of dorsal (medial) margin of external naris (left side) |
| 34 | Lateral extreme of frontonasal contact (left side) |
| 35 | Lateral extreme of posterior margin of basisphenoid (right side) |
| 36 | Lateral extreme of posterior margin of basisphenoid (left side) |
| 37 | Dorsal contact between premaxilla and jugal bar (right side) |
| 38 | Dorsal contact between premaxilla and jugal bar (left side) |

**Table S4. Semi-landmark curve guide (see anatomical landmark guide for initial and terminal landmarks).** Adapted from Mitchell et al., 2021.

| Curve | Number of sliding semi-landmarks | Initial Landmark | Terminal Landmark | Description |
| --- | --- | --- | --- | --- |
| 1 | 30 | 1 | 2 | Midline of the palate |
| 2 | 20 | 1 | 3 | Lateral margin of palate |
| 3 | 10 | 4 | 3 | Lateral margin of premaxilla |
| 4 | 15 | 3 | 37 | Perimeter of jugal bar |
| 5 | 15 | 37 | 6 | Anterior margin of the antorbital fenestra |
| 6 | 20 | 6 | 8 | Lateral margin of the orbit |
| 7 | 10 | 4 | 5 | Midline of the dorsal side of rostrum |
| 8 | 10 | 5 | 9 | Midline of the dorsal side of cranial vault |
| 9 | 10 | 9 | 10 | Midline of supraoccipital |
| 10 | 10 | 10 | 11 | Lateral margin of foramen magnum |
| 11 | 10 | 11 | 12 | Lateral margin of occipital condyle |
| 12 | 20 | 9 | 13 | Lateral margin of occipital complex |
| 13 | 5 | 13 | 12 | Midline of suboccipital region |
| 14 | 30 | 7 | 3 | Posterolateral margin of palate |
| 15 | 20 | 13 | 14 | Midline of basisphenoid |
| 16 | 10 | 15 | 16 | Posterior margin of articular surface of quadrate |
| 17 | 10 | 16 | 15 | Anterior margin of articular surface of quadrate |
| 18 | 10 | 17 | 7 | Medial edge of pterygoid |
| 19 | 10 | 7 | 18 | Lateral edge of pterygoid |
| 20 | 10 | 18 | 17 | Contact between pterygoid and quadrate |
| 21 | 10 | 19 | 20 | Ventrolateral margin of naris |
| 22 | 10 | 19 | 21 | Dorsomedial margin of naris |
| 23 | 10 | 5 | 22 | Craniofacial hinge/frontonasal suture |
| 24 | 20 | 14 | 35 | Lateral margin of basisphenoid |

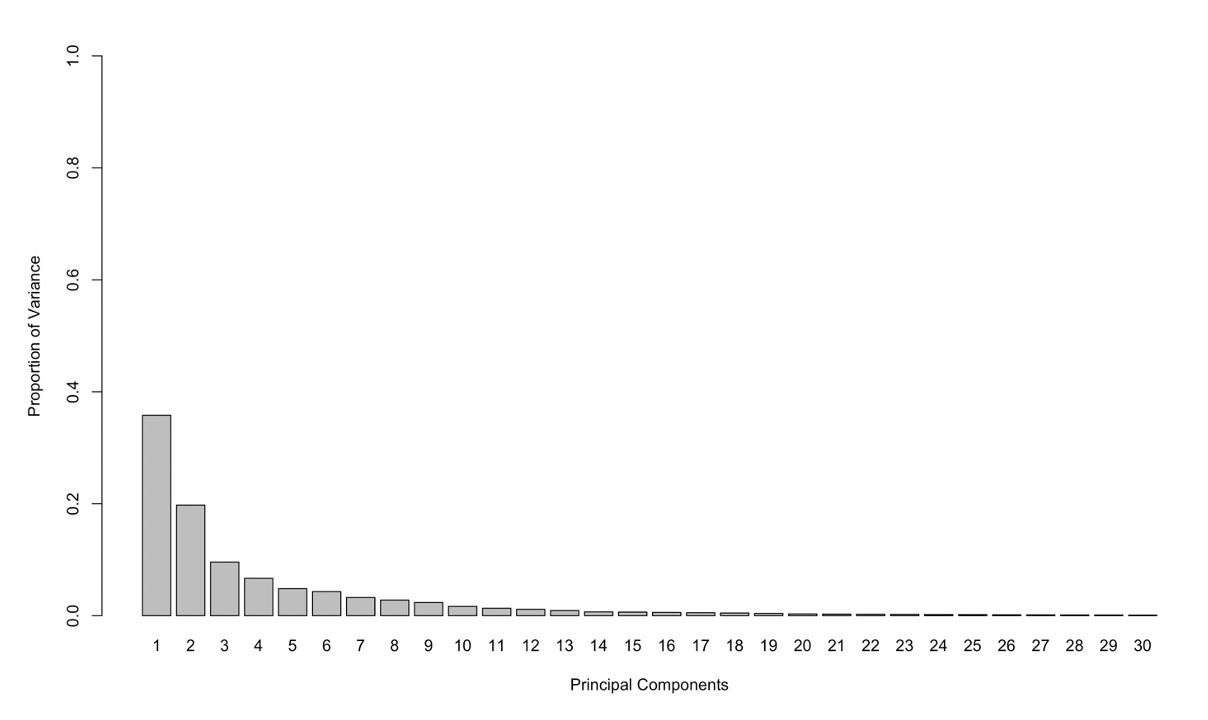
**Figure S1. Proportional Variance of principal components of the full dataset.**

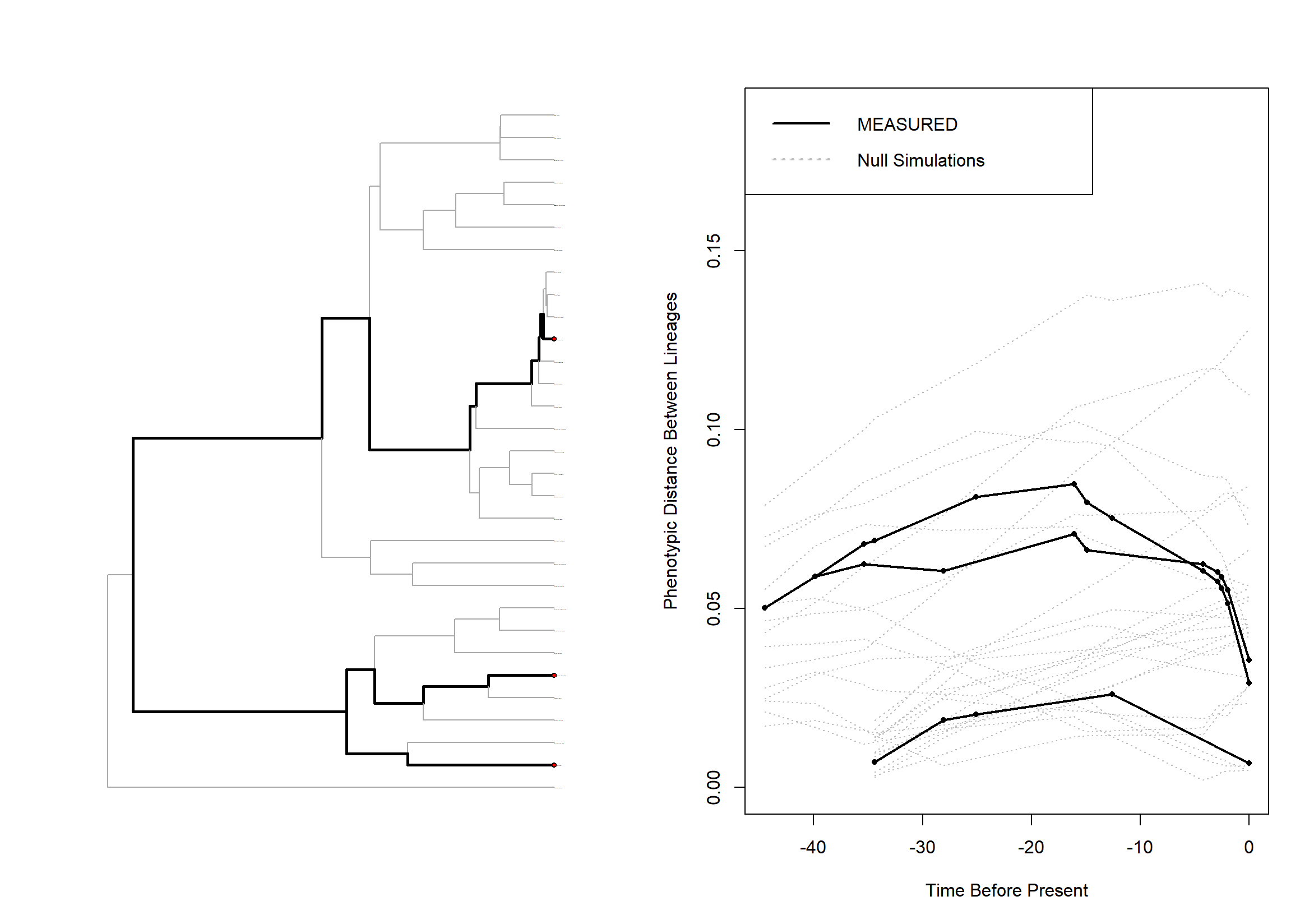

**Figure S2: Convergence plot for three gulper taxa (*Vultur gryphus, Gymnogyps californianus, Gyps tenuirostris*; Ct_1_ = 0.63, *p* = 0.00).**

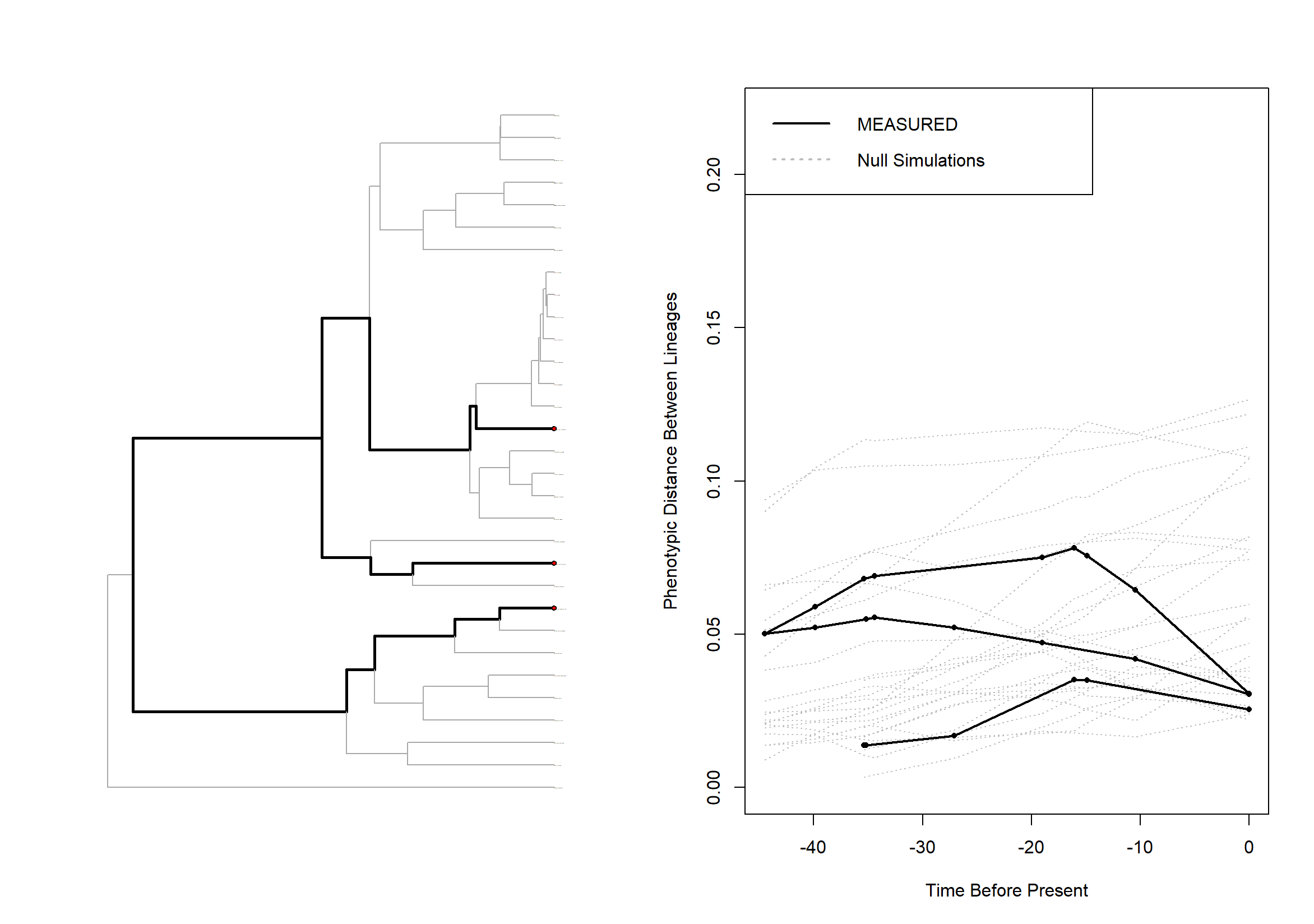

**Figure S3: Convergence plot for three scrapper taxa (*Necrosyrtes monachus, Neophron percnopterus, Cathartes melambratus*; Ct_1_ = 0.45, *p* = 0.00)**

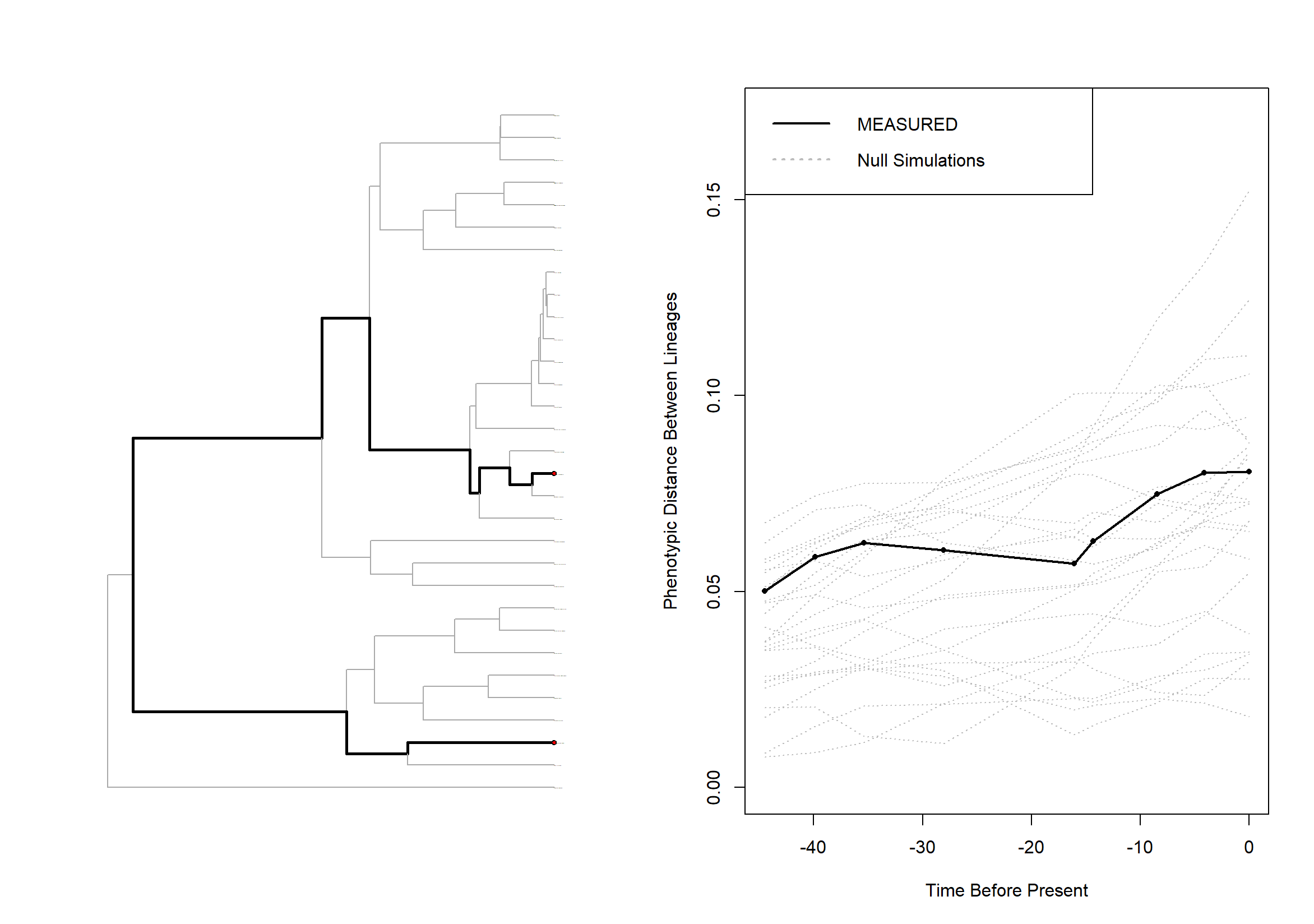

**Figure S4: Convergence plot for two ripper taxa (*Sarcoramphus papa, Torgos tracheliotos;* Ct_1_ = -0.004, *p* = 0.38)**

**Table S6. Summary of phylogenetic signal (*K_mult_*) across all datasets.**

| Shape Data | *K_mult_* | Effect Size | P-value |
| --- | --- | --- | --- |
| All species | 0.33 | 7.95 | 0.001 |
| All species, allometry-corrected | 0.26 | 5.19 | 0.001 |
| Vultures-only | 0.26 | 5.13 | 0.001 |
| Vultures-only, allometry-corrected | 0.21 | 3.65 | 0.001 |
| Feeding Type-only | 0.26 | 4.25 | 0.001 |
| Feeding Type-only, allometry-corrected | 0.19 | 3.07 | 0.001 |

**Table S7. Summary of analyses on all Procrustes-aligned subsets of the data.**

| Statistical Test | All taxa | | | | Vultures only | | | | Known feeding ecology only | | | |
| --- | --- | --- | --- | --- | --- | --- | --- | --- | --- | --- | --- | --- |
|  | **R^2^** | **F** | **Z** | ***p*** | **R^2^** | **F** | **Z** | ***p*** | **R^2^** | **F** | **Z** | ***p*** |
| Allometry, raw shape data | 0.185 | 6.59 | 3.63 | **0.001** | 0.188 | 4.63 | 2.90 | **0.001** | 0.207 | 4.69 | 2.81 | **0.001** |
| Phylogenetic Allometry | 0.050 | 1.54 | 1.46 | 0.073 | 0.033 | 0.68 | -0.27 | 0.587 | 0.042 | 0.78 | 0.06 | 0.51 |
| MANOVA Vulture/Non-vulture | 0.158 | 5.44 | 3.27 | **0.001** | NA | NA | NA | NA | NA | NA | NA | NA |
| Phylogenetic MANOVA Vulture/Non-vulture | 0.013 | 0.381 | -1.23 | 0.894 | NA | NA | NA | NA | NA | NA | NA | NA |
| Feeding Type MANOVA | 0.458 | 7.17 | 4.79 | **0.001** | 0.458 | 7.17 | 4.79 | **0.001** | 0.458 | 7.17 | 4.79 | **0.001** |
| Allometry-corrected Feeding Type | 0.389 | 5.41 | 3.72 | **0.001** | 0.357 | 4.72 | 3.46 | **0.001** | 0.356 | 4.70 | 3.44 | **0.001** |
| Allometry*Feeding Type interaction | 0.068 | 1.10 | 0.382 | 0.354 | 0.068 | 1.10 | 0.38 | 0.354 | 0.068 | 1.10 | 0.38 | 0.354 |
| Family*Feeding Type interaction | 0.089 | 2.29 | 2.05 | **0.028** | 0.089 | 2.29 | 2.04 | **0.028** | 0.088 | 2.29 | 2.04 | **0.028** |
| Phylogenetic MANOVA Feeding Type | 0.090 | 0.832 | 0.160 | 0.436 | 0.085 | 0.79 | 0.05 | 0.483 | 0.085 | 0.79 | 0.05 | 0.482 |
| Phylogenetic Feeding Type*Allometry interaction | 0.116 | 1.05 | -0.015 | 0.524 | 0.122 | 1.11 | 0.08 | 0.478 | NA | NA | NA | NA |

**Table S8. Flexible discriminant analysis – vulture feeding type.** Predicted group assignment for vulture feeding type, implemented with *mvMorph* (Clavel et al., 2015).

| Species | Gulper | Ripper | Scrapper |
| --- | --- | --- | --- |
| *Breagyps clarki* | 1 | 0 | 0 |
| *Gypaetus barbatus* | 0 | 0 | 1 |
| *Gypohierax angolensis* | 0 | 0 | 1 |
| *Hieraaetus moorei* | 0 | 0 | 1 |

**Table S9. Flexible discriminant analysis – feeding group.** Predictions for feeding group with added ‘raptor’ category, implemented with *mvMorph* (Clavel et al., 2015).

| Species | Gulper | Ripper | Scrapper | Raptor |
| --- | --- | --- | --- | --- |
| *Breagyps clarki* | 1 | 0 | 0 | 0 |
| *Gypaetus barbatus* | 0 | 0 | 0 | 1 |
| *Gypohierax angolensis* | 0 | 0 | 1 | 0 |
| *Hieraaetus moorei* | 0 | 0 | 0 | 1 |

**Table S10. Flexible discriminant analysis – vulture/non-vulture.** Predictions for bird type, implemented with *mvMorph* (Clavel et al., 2015).

| Species | Non-Vulture Raptor | Vulture |
| --- | --- | --- |
| *Breagyps clarki* | 0 | 1 |
| *Gypaetus barbatus* | 0 | 1 |
| *Gypohierax angolensis* | 0 | 1 |
| *Hieraaetus moorei* | 1 | 0 |

**Table S11. Flexible discriminant analysis – vulture/non-vulture.** Predictions for bird type, implemented with *mvMorph* (Clavel et al., 2015)..

| Species | Non-Vulture Raptor | Vulture |
| --- | --- | --- |
| *Breagyps clarki* | 0 | 1 |
| *Hieraaetus moorei* | 1 | 0 |
